# The desmin mutation R349P increases contractility and fragility of stem cell-generated muscle micro-tissues

**DOI:** 10.1101/2021.07.23.453481

**Authors:** Marina Spörrer, Delf Kah, Richard C. Gerum, Barbara Reischl, Danyil Huraskin, Claire A. Dessalles, Werner Schneider, Wolfgang H. Goldmann, Harald Herrmann, Ingo Thievessen, Christoph S. Clemen, Oliver Friedrich, Said Hashemolhosseini, Rolf Schröder, Ben Fabry

**Author notes:** These authors contributed equally to this work (Marina Spörrer, Delf Kah).

## Abstract

Desminopathies comprise hereditary myopathies and cardiomyopathies caused by mutations in the intermediate filament protein desmin that lead to severe and often lethal degeneration of striated muscle tissue. Animal and single cell studies hinted that this degeneration process is associated with massive ultrastructural defects correlating with increased susceptibility of the muscle to acute mechanical stress. The underlying mechanism of mechanical susceptibility, and how muscle degeneration develops over time, however, has remained elusive. Here, we investigated the effect of a desmin mutation on the formation, differentiation, and contractile function of *in vitro*-engineered three-dimensional micro-tissues grown from muscle stem cells (satellite cells) isolated from heterozygous R349P desmin knock-in mice. Micro-tissues grown from desmin-mutated cells exhibited spontaneous unsynchronized contractions, higher contractile forces in response to electrical stimulation, and faster force recovery compared to tissues grown from wild-type cells. Within one week of culture, the majority of R349P desmin-mutated tissues disintegrated, whereas wild-type tissues remained intact over at least three weeks. Moreover, under tetanic stimulation lasting less than five seconds, desmin-mutated tissues partially or completely ruptured, whereas wild-type tissues did not display signs of damage. Our results demonstrate that the progressive degeneration of desmin-mutated micro-tissues is closely linked to extracellular matrix fiber breakage associated with increased contractile forces and unevenly distributed tensile stress. This suggests that the age-related degeneration of skeletal and cardiac muscle in patients suffering from desminopathies may be similarly exacerbated by mechanical damage from high-intensity muscle contractions. We conclude that micro-tissues may provide a valuable tool for studying the organization of myocytes and the pathogenic mechanisms of myopathies.

## Introduction

Desmin is the primary intermediate filament in skeletal, cardiac, and smooth muscle tissue (1–4). Desmin filaments form a scaffold around the myofibrillar Z-discs, connecting the contractile apparatus to the subsarcolemmal cytoskeleton, myo-nuclei, mitochondria, and other organelles. Thus, the desmin cytoskeletal network plays a central role in myofibrillar integrity and force transmission (1, 2, 4, 5). It is therefore not surprising that a number of desmin mutations lead to disturbed filament formation in particular under mechanical stress, impaired force transmission, and ultimately to myofibrillar degeneration (2, 4, 6, 7). A meta-analysis of 159 patients with desmin gene mutations reported that more than 70% of the patients developed myopathies, and 50% developed cardiomyopathies. Desminopathies are rare in humans (less than 50 cases per 100,000 individuals (4)), but are associated with severely reduced quality of life and life expectancy (2).

Over the last decades, our understanding of desminopathies has been greatly improved by the use of mouse models. Numerous studies have shown that mice either lacking wild-type desmin or expressing mutant desmin develop disorganized muscle fibers (8, 9), desmin-positive aggregates (10), skeletal muscle weakness (10, 11), increased muscle vulnerability during contraction (11), and mitochondrial dysfunction (8, 12, 13). Other studies confirmed these results in zebrafish. Larvae carrying a desmin mutation were smaller than wild-type animals, their swimming activity was reduced, and their muscles showed structural disorganization (14). Furthermore, disruption of the excitation-contraction (EC) coupling machinery and abnormal subcellular localization of ryanodine receptors (RyR) were detected in desmin-negative zebrafish (15).

Studies on NIH-3T3 fibroblasts transfected with the most common desmin mutation in humans, R350P, showed disrupted intermediate filament (IF) networks as well as cytoplasmic protein aggregates, which are in agreement with findings in animal studies (16). However, these and other single cell studies have not provided insight into mechanisms that explain how cytoskeletal network disorganization and the presence of desmin-positive aggregates lead to muscle dysfunction at the tissue level. Moreover, a follow-up study found that several other desmin mutations that cause diseases in humans and animals did not alter filament formation at the single cell level (17).

To explore potential mechanisms that link desmin-related structural aberrations at the cellular level to the increased mechanical vulnerability and loss-of-function at the level of muscle tissue, we generated bioartificial three-dimensional (3-D) micro-tissues from desmin-mutated cells. We reasoned that such 3-D micro-tissues may better reproduce the *in vivo* situation compared to classical single cell studies, while at the same time being easier to handle and providing higher measurement throughput than whole muscle tissue or muscle fibers dissected from animals.

Our approach is based on a previously established micro-tissue system, where muscle cells are mixed with unpolymerized collagen-I and Matrigel in a small well containing two flexible pillars (18–20). Upon polymerization of matrix proteins into a fibrillar network, the embedded cells begin to adhere, spread, and generate contractile forces. These forces compact the matrix network fibers, which wrap around and align between the flexible pillars. The cells in turn align with the parallel matrix fiber bundles, fuse, and over time form a 3-D contractile tissue. The force generation of this tissue can be measured by the deflection of the flexible pillars. 3-D micro-tissues grown from skeletal muscle cell sources can thus be used as a platform for studying physiological and mechanical properties of muscles (20, 21), for drug testing (22, 23), and for disease modeling (23–25).

As the cell source for generating micro-tissues, we used skeletal muscle stem cells (satellite cells) from wild-type mice and from mice carrying an R349P desmin mutation, which is the ortholog of the R350P mutation in humans. In addition, we added electrical stimulation electrodes to the 3-D culture system to measure both the static forces (baseline tone) and the active forces (contractile force in response to electrical stimulation) of the muscle micro-tissues. We found that micro-tissues grown from desmin-mutated satellite cells exhibited decreased static forces but increased active contractile forces compared to wild-type tissues, indicating pronounced hypercontractility. Furthermore, mutated tissues showed less organized fiber structures and were prone to tissue disintegration in response to tetanic electrical stimulation.

## Materials and methods

### Mice Strains, Skeletal muscle cell extractions and culturing conditions

The R349P desmin knock-in mouse model (B6J.129Sv-*Des*^tm1.1Ccrs^/Cscl MGI:5708562; hereafter “Des^R349P^”) was generated as previously described (10). Mouse experiments were performed following animal welfare laws and approved by the responsible local committee (animal protection officer, Sachgebiet Tierschutzangelegenheiten, FAU Erlangen-Nürnberg, reference numbers: I/39/EE006, TS-07/11). Mice were housed in cages at 22 ± 1 °C and relative humidity of 50–60% on a 12 h light/dark cycle. Water and food were provided ad libitum.

We prepared satellite cells from two wild-type (wt) BL6 mice and two heterozygous Des^R349P^ (hereafter “Des^wt/R349P^”) BL6 mice (age 2-3 months). Primary muscle satellite cells were extracted and pooled from 1-2 g of striated skeletal muscle tissue per mouse using the MACS Satellite Cell Isolation Kit (Miltenyi Biotech) according to the manufacturer’s instructions (26). Cells were seeded at a density of about 1,000 cells/cm^2^ in culture dishes (353003, Corning) coated with Matrigel. For this purpose, 1% Matrigel (Corning) in culture medium was applied to the dishes in a thin layer (10 µl of the solution per culture dish was spread with a cell scraper) and then incubated for two hours at 37°C and 5% CO_2_. The culture dishes could be used directly or stored with the addition of medium at incubation conditions. Culture medium consisted of Dulbecco’s Modified Eagle Medium (DMEM, Gibco, 41966) supplemented with 40% (v/v) Ham’s F10 nutrient mixture (Gibco), 20% (v/v) fetal calf serum, 1% (v/v) penicillin-streptomycin (Thermo Fisher, 15070, containing 10,000 units/ml of penicillin and 10,000 µg/ml of streptomycin) and 5 ng/ml recombinant human fibroblast growth factor (G5071, Promega).

### Fabrication of 3-D micro-tissues

The protocol for fabricating 3-D muscle micro-tissues was adapted from (18). Tissues were grown in PDMS devices containing 18 small wells with dimensions of 3.6 × 1.8 × 2 mm (l × w × h) with two flexible pillars (diameter 500 µm, 2.5 mm height, 1.8 mm center-to-center distance) (Fig. 1A). The PDMS chambers were coated with a solution of 1% Pluronic F-127 acid (Sigma-Aldrich) in water overnight to prevent matrix proteins from sticking to the walls of the PDMS chambers. After aspirating the coating solution, 6 µl of an unpolymerized ice-cold collagen-I/ Matrigel solution was pipetted into each well. 1 ml of solution contained 100 µl of 2 mg/ml Collagen R (collagen I from rat tail veins, Matrix Bioscience, 50301), 77 µl of 5.2 mg/ml Collagen G (collagen I from bovine calf skin, Matrix Bioscience, 50104), 100 µl of 15 mg/ml Matrigel (Corning), 35.1 µl NaHCO_3_ (23 mg/ml, Sigma-Aldrich), 35.1 µl 10x DMEM (Seraglob), 4.2 µl NaOH (1M) and 649 µl of dilution medium that consisted of one volume part NaHCO_3_ (23 mg/ml), one volume part 10x DMEM, and eight volume parts distilled H_2_O, adjusted to pH 10 using NaOH. The PDMS devices were centrifuged (1 min at 300g) and incubated at 37°C and 5% CO_2_ for 1 h, during which the collagen-I/Matrigel solution polymerized. Next, a second layer of 6 µl unpolymerized ice-cold collagen-I/Matrigel solution mixed with 5,000 cells was pipetted on top of the polymerized bottom layer in each PDMS chamber, allowing it to polymerize for 1 hour at 37°C and 5% CO_2_. Then, 1 ml of culture medium was carefully pipetted on top of the wells of each device.

**Figure 1.**
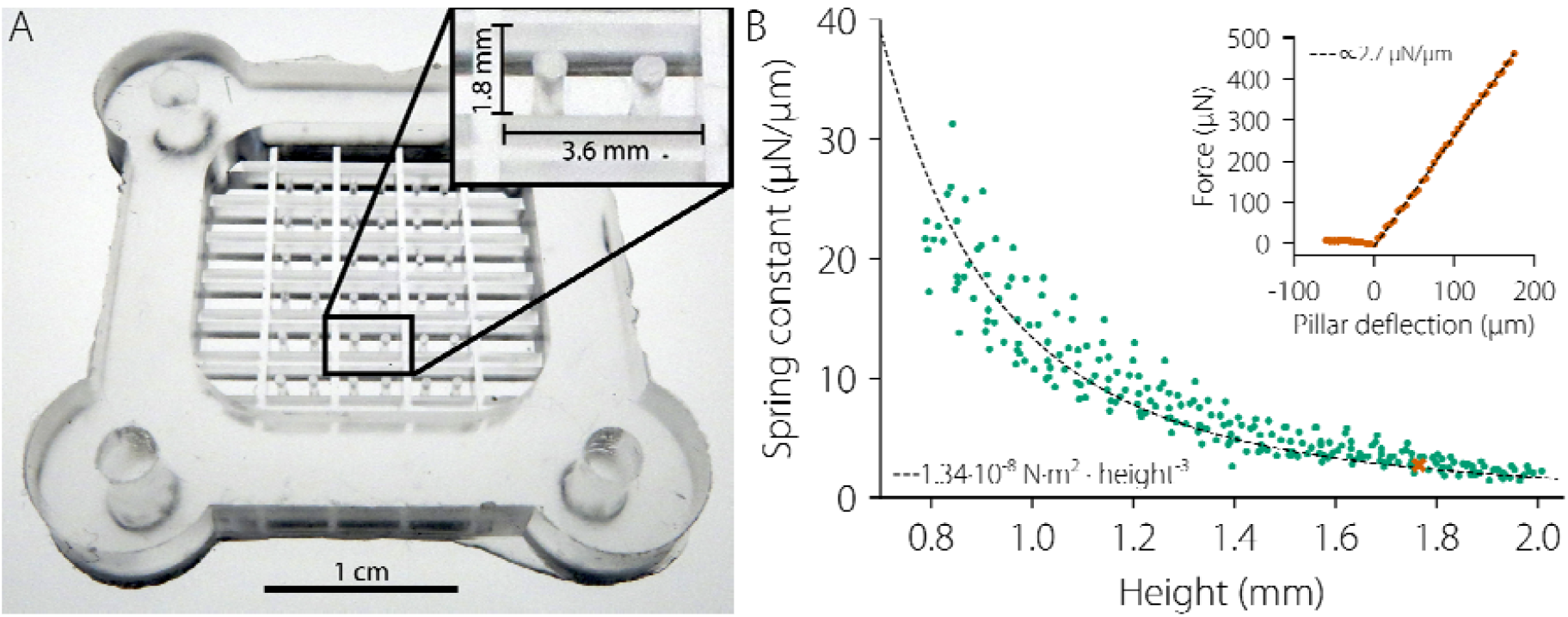
**A**. PDMS device for the culture of muscle micro-tissues. **B**. Calibration curve for determining PDMS pillar spring constants. The spring constants of PDMS pillars were determined at different heights with respect to the bottom of the PDMS well by measuring the force required for a defined bending (pillar deflection). A representative measurement of the force-deflection relationship at a height of 1.75 mm is shown in the inset. The spring constant is then computed from the slope of the force-deflection relationship. Green data points indicate the spring constants of individual measurements at different heights. The dotted line shows the prediction of the Euler-Bernoulli beam theory, according to which the effective spring constant is proportional to the negative third power of the height.

Upon polymerization, the collagen-I/Matrigel solution forms a fiber network to which the skeletal muscle cells adhere, spread, and apply forces, causing a collective re-modelling of the network into a dense tissue with parallel bundles of fibers that span between the two pillars (Fig. 2A). This process of tissue formation is completed typically within 12 h after seeding. 24 h after seeding, the culture medium was exchanged for a differentiation medium, which was changed every day. Differentiation medium consisted of DMEM supplemented with 2% horse serum, 1% penicillin-streptomycin and 0.5% insulin-transferrin-selenium (Gibco, 41400, consisting of 1 g/l insulin, 0.55 g/l transferrin, and 670 ng/l sodium selenite).

**Figure 2.**
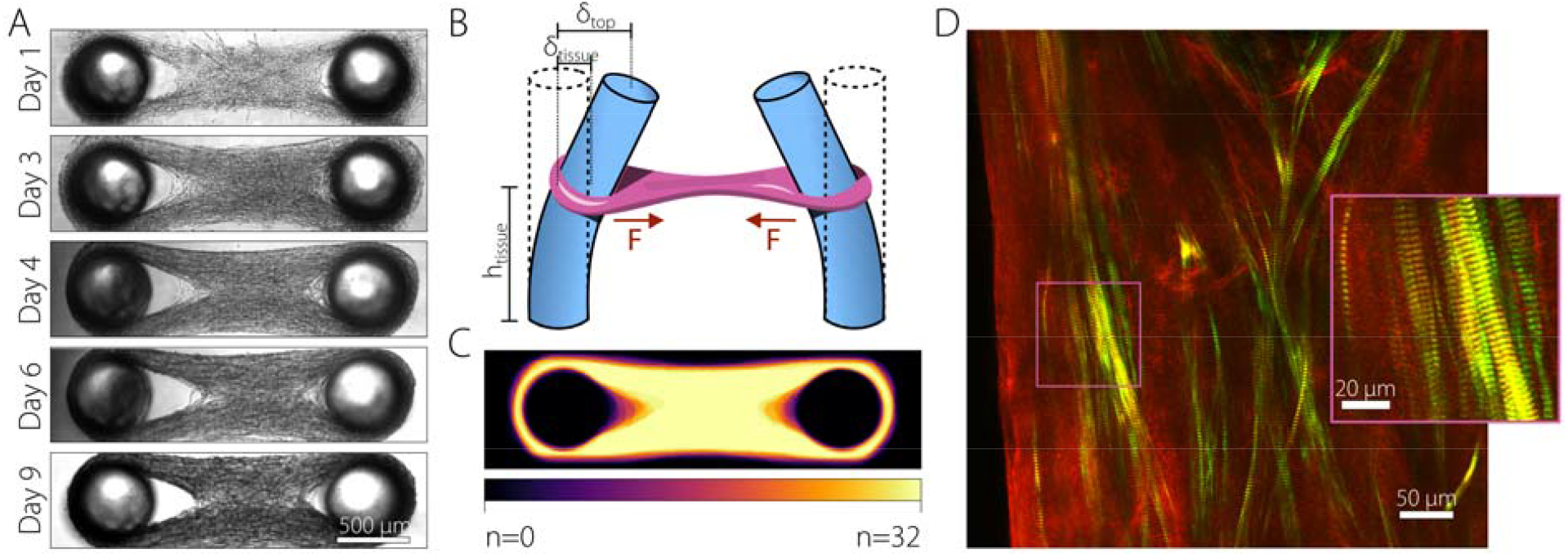
Morphology of wild-type satellite tissues. **A**. Bright-field images of tissue compaction from day 1 to day 9. **B**. Schematic of pillars and tissue during force generation. **C**. Overlay heatmap of the morphology of 32 tissues on day 4 of differentiation. **D**. SHG multiphoton confocal image of a tissue on day 21 after differentiation. Green=myosin, red=collagen.

### Evaluation of contractile force

The axial contractile force *F* of a micro-tissue was measured from the pillar deflection *δ* using Hooke’s law

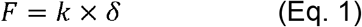

where *k* is the spring constant of a PDMS pillar at the height of the tissue above the pillar base *h*_*tissue*_ (Fig. 2B). *h*_*tissue*_ was measured by manually focusing a bright-field microscope to the tissue regions near the pillars. *k* was determined from the force-bending relationship measured at different heights above the pillar base, using a motorized micromanipulator (InjectMan NI2, Eppendorf AG) and a laboratory scale (Practum 64-1S, Sartorius). The force (*F*) versus pillar deflection (*δ*) relationship at different heights (*h*) was then fitted to the Euler-Bernoulli beam equation: *F(h)* = *k*_*0*_/*δ*(*h/h*_*0*_)^3^, where *h*_*0*_ is a reference height that we arbitrarily set to 1 mm, and *k*_*0*_ is the spring constant of the pillars at a height of *h*_*0*_. For the devices used in this study, *k*_*0*_ was around 10 µN/µm. The pillar spring constant *k* at different heights can then be computed according to *k = k*_*0*_(*h*/*h*_*0*_)^3^ (Fig. 1B).

For small contractile forces of the tissues, the pillar deflection *δ* at the tissue height is on the order of a few µm. For higher accuracy, we therefore measured the deflection *δ*_*top*_ at the top of the pillar, where the deflection is larger than at the height of the tissue *h*_*tissue*_ (Fig. 2B). *δ* can then be computed using the Euler-Bernoulli beam theory according to

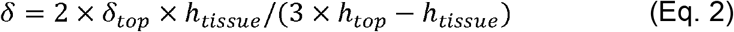

where *h*_*top*_ is the total pillar length (2.5 mm in our setup).

In the following, we distinguish between the static and the active force. Static force is defined as the pillar bending force when the myocytes are not electrically paced. It is typically measured as the lowest force between two pacing pulses at least 1 s apart. The active force is defined as the maximum contractile force exerted by the tissues above the static force in response to electrical stimulation.

### Electrical stimulation

The micro-tissues were electrically stimulated via four vertical graphite electrodes (Faber Castell pencil lead type “F” with 0.5 mm diameter) that dipped approximately 3 mm into the cell culture medium. The electrodes were rectangularly arranged in two pairs, with a distance of 9 mm between the two electrodes with equal polarity, and a distance of 18 mm between the pairs of opposite polarity. The stimulation signal was a biphasic square wave pulse consisting of a positive pulse with a field strength of 10 V/cm and 5 ms duration, followed by a negative pulse with a field strength of −10 V/cm and 5 ms duration. The pulse repetition rate was either 1 Hz for single pulse mode or 100 Hz for tetanic stimulation.

### Chemical stimulation

For chemical stimulation, we “skinned” the micro-tissues with 10 mg/ml saponin in an EGTA-buffered calcium-free salt solution for three minutes. The calcium-free salt solution with an osmolality of 161 mOsm/kg as measured with an osmometer (Osmomat3000, Gonotec, Berlin) contained 30 mM Hepes, 6.25 mM Mg(OH)_2_, 30 mM EGTA, 8 mM Na_2_ATP, and 10 mM Na_2_CP, and the pH was adjusted to 7.0 using 1 M KOH. Then, we washed the sample in a calcium-free salt solution and measured the contractile forces of five micro-tissues. We repeated the measurement eleven times and replaced the medium with a salt solution of gradually increasing calcium concentration before each measurement. The negative decadic logarithm of the calcium concentration (pCa) ranged from 6.74 (1.82 × 10^−7^ M) to 4.92 (1.19 × 10^−5^ M). pCa-values and contractile forces were plotted and fitted to a generalized Hill equation of the form,

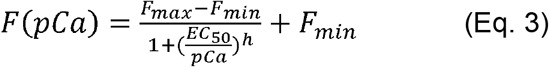

where *F*_*min*_ and *F*_*max*_ are the minimum and maximum saturation levels of the Hill curve, *EC*_*50*_ is the half maximum effective concentration, and *h* is the Hill coefficient.

### Imaging

Bright-field images were taken with a charge-coupled device camera (Hamamatsu, ORCA ER). Second harmonic generation (SHG) confocal images were taken using a multiphoton microscope (TriMScope II, LaVision BioTec) as described previously (27). In brief, the SHG signals of myosin-II and collagen-I were excited with a Ti:Sa laser (Charmeleon Vision II, Coherent) with a power of approximately 16 mW and a pulse duration of approximately 150 fs. The backward scattered signal (405 ± 10 nm) was used for detection of fibrillar collagen. The forward scattered signal (405 ± 10 nm) was used for the detection of myosin II. The SHG signals were detected using a non-descanned transmission photomultiplier tube (H 7422-40 LV 5 M, Hamamatsu Photonics). Images were taken with a 25x dipping objective (Leica, HC Fluotar NA 0.95) with a line average of three and at a frequency of 1,000 Hz. The resolution was 1,024 × 1,024 pixels (481.3 × 481.3 µm), and the z-stacks had a step-size of 0.5 µm.

### Statistical Analysis

Values are expressed as mean ± standard deviation (SD) from measurements of different micro-tissues. The statistical significance of the differences between groups (wild-type versus desmin-mutated) was assessed using a Student’s two-tailed t-test assuming unequal variance. Boxplots indicate median values, lower to upper inter quartile values, 1.5 interquartile ranges as well as individual force measurements (as a scatter plot).

## Results

### Formation of micro-tissues

We mixed satellite wild-type cells with unpolymerized collagen-I/Matrigel solution and seeded them in PDMS chambers containing two pillars. After the collagen-I/Matrigel solution polymerized into a fiber network, the cells started to adhere, spread, and collectively exert contractile forces. In response, the network fibers and cells aligned between the pillars. Within 24 hours after seeding, the cells had compacted the fiber network into a dog bone-shaped micro-tissue (Fig. 2A, “Day 1”). We then replaced the culture medium with a differentiation medium supplemented with insulin, transferrin, and selenium (ITS). Over the following days, the micro-tissues increased in width and density (Fig. 2A). Typically, four days after seeding, the pillars started to bend inwards, indicating that the micro-tissues generated measurable contractile forces (Fig. 2A, B). The morphology of wild-type micro-tissues was highly reproducible (Fig. 2C). A heat map of overlaid bright-field images from 32 different micro-tissues after four days of differentiation (five days after seeding) showed only slight differences in tissue morphology around the fusion region of the two tissue branches that wrap around the pillars. Second harmonic generation confocal images of collagen and myosin-II showed highly elongated, well aligned cells with the typical striation pattern of differentiated myofibers (Fig. 2D).

### Micro-tissues from wild-type satellite cells recapitulate the contractile behavior of intact muscle

In tissues from wild-type satellite cells, we investigated static and active forces over 20 days of differentiation. Active force was measured as the change in force in response to electrical stimulation (± 10 V/cm bipolar 5 ms pulses at 1 Hz), and the static force was measured as the minimum force between two electrical stimulation pulses (Fig. 3A). We compared static and active forces in tissues grown from three different cell passages (Fig. 3B, D). The static force was around 60-70 µN for tissues from all three passages and remained approximately constant over time. The active force was around 50 µN on average, but exhibited a slightly higher variability between cell passages. Active and static forces slowly decreased over time in tissues from passage 1, whereas they slowly increased over time in tissues from passage 3 and 4. Importantly, active forces remained substantial in all tissues over a duration of 3 weeks.

**Figure 3.**
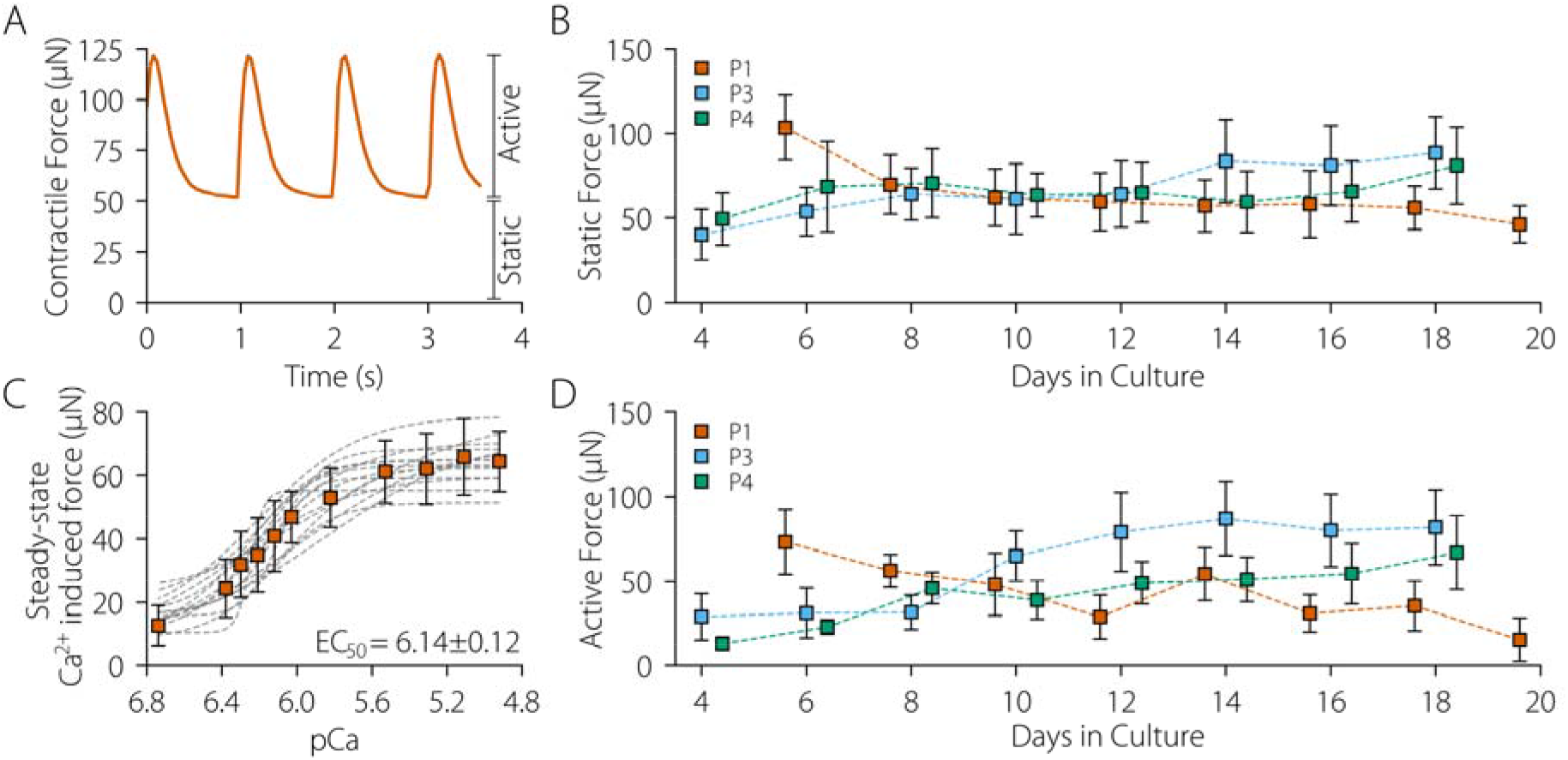
Static and active contractile forces in wild-type satellite cell tissues. **A**. Static force and active contractile force in response to electrical stimulation in a representative wild-type tissue (day 10 of differentiation) **B**. Static force (mean ± SD) from day 6 to day 20 of differentiation in tissues derived from three different cell passages, with number of tissues n(P1) = 18, n(P3) = 21, and n(P4) = 17. **C**. Dependence of the contractile force (mean ± SD of 15 tissues) on calcium concentration in “skinned” micro-tissues. Gray lines represent the contractile response of individual micro-tissues. **D**. Same as in Fig. 3B for active forces.

To compare the force that the tissues generated in response to electrical versus chemical stimulation, we permeabilized the cell membrane with saponin and gradually increased the Ca^2+^ concentration in the medium from 6.74 to 4.92 (pCa). The total contractile steady-state force showed the typical sigmoidal dependence on pCa as reported for skinned single muscle fibers (28–30) (Fig. 3C, S1). The pCa-concentration for half-maximum forces was 6.14 ± 0.12, and the maximum force was 65.3 ± 7.4 µN, similar to the active force in response to electrical excitation in these tissues (cell passage 14, after 9 days of differentiation).

### R349P mutant desmin induces structural instability and altered contractile behavior

We next tested the morphology and contractile performance of micro-tissues generated using satellite cells from *Des*^*wt/*R349P^ mice. These tissues compacted similarly fast and were morphologically comparable to wild-type tissues at early time points. However, after around day 3 of differentiation, *Des*^*wt/*R349P^ tissues started to disintegrate. Individual muscle fibers ruptured at their thinnest points near the pillar, resulting in progressive cell aggregation in the central region of the tissue (Fig. 4A). A heat map of bright-field images from 21 different micro-tissues after four days of differentiation showed a pronounced variability in tissue morphology (Fig. 4B). Notably, on day 6, about two thirds of *Des*^*wt/*R349P^ tissues had ruptured, whereas no wild-type tissue had ruptured at this time point.

**Figure 4.**
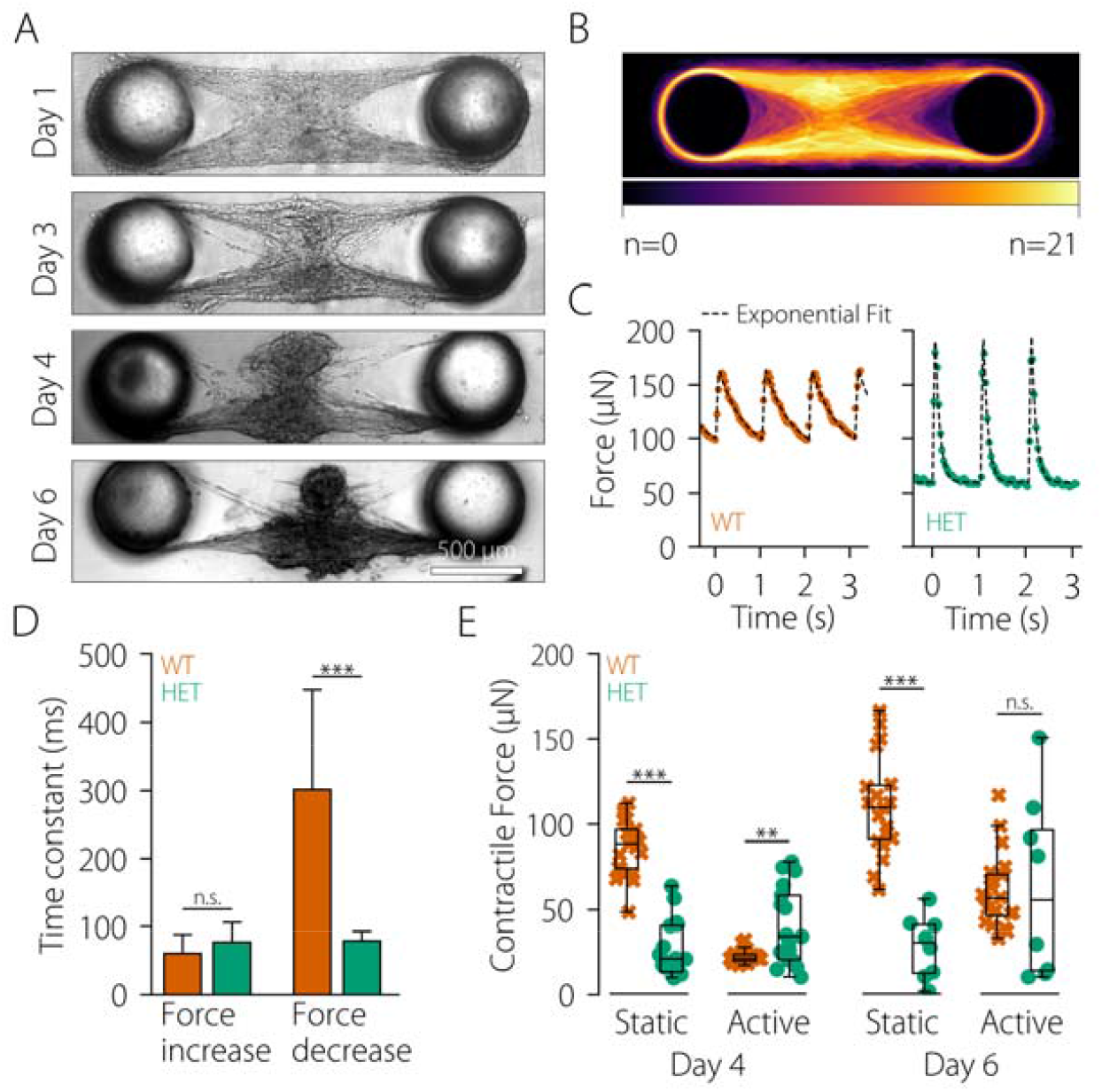
Morphology and force generation in tissues with a heterozygous desmin mutation. **A**. Bright-field images of tissue compaction from day 1 to day 6 of differentiation. **B**. Overlay heatmap of the morphology of 21 tissues on day 4 of differentiation. **C**. Force vs. time in a typical wild-type (orange) and Des^wt/R349P^ (green) tissue after 6 days of differentiation. Dots represent measurement points and dotted lines represent the superposition of two exponential functions with a short time constant for phases with increasing forces and a longer time constant for phases with decreasing forces. **D**. Time constants for force increase and force decrease (mean ± SD) in wild-type (orange, n=19) and Des^wt/R349P^ (green, n=9) tissues after 6 days of differentiation. **E**. Active and static force of wild-type (orange) and Des^wt/R349P^ (green) tissues on day 4 and day 6 of differentiation. *p ≤ 0.05, **p ≤ 0.01, ***p ≤ 0.001, n.s., not significant (p > 0.05).

We also observed substantial differences in the contractile behavior of wild-type versus desmin-mutated tissues. The duration of a single pulse contraction (twitch) was considerably shorter in *Des*^*wt/*R349P^ tissues (Fig. 4C). From a fit of two superimposed exponential functions to the force-time trace, we found that the time constant for the force increase in response to an electrical pulse was similar (61 ± 28 ms for wild-type and 77 ± 30 ms for mutated tissues, mean ± SD). However, the time constant for force relaxation was 4-times lower in the mutated tissues (79 ± 15 ms) compared to wild-type tissues (301 ± 147 ms). In addition, on day 4 of differentiation, *Des*^*wt/*R349P^ tissues showed lower static contractile forces (26.7 ± 16.5 µN) compared to wild-type tissues (40.3 ± 23.0 µN) (Fig. 4E). These differences in static force were even more pronounced on day 6 of differentiation with 28.3 ± 18.5 µN in mutated tissues compared to 110.6 ± 29.4 µN in wild-type tissues. By contrast, on day 4 of differentiation, *Des*^*wt/*R349P^ tissues generated significantly (p < 0.01) higher active forces (40.3 ± 23.0 µN) compared to wild-type tissue (22.5 ± 4.1 µN).

The relative variability in the active force between the individual tissues, as quantified by the coefficient of variance (COV), was also considerably larger in mutated tissues (0.57) compared to wild-type tissues (0.18). The differences in active force diminished after day 6 of differentiation (62.8 ± 53.2 µN in *Des*^*wt/*R349P^ tissues versus 60.8 ± 21.9 µN in wild-type tissues), but the variability of active forces between individual tissues remained larger in *Des*^*wt/*R349P^ tissues (0.85) compared to wild-type tissues (0.36). Together, these data demonstrate that heterozygous knock-in of *Des*^R349P^ is associated with an increased active contractile force generation, but with reduced static tissue forces and impaired tissue integrity.

### Tetanic stimulation causes rupture of desmin-mutated tissues

We next tested the hypothesis that the progressive rupture of *Des*^*wt/*R349P^ tissues was driven by larger contractile forces. We reasoned that in combination with thinner tissue fibers that wrap around the pillars, larger forces may lead to considerably higher mechanical stress and ultimately tissue failure. To test this idea, we applied tetanic electrical stimulation (100 Hz pulse trains for several seconds), which led to 1.75 fold higher active force generation compared to single twitch stimulation, both in *Des*^*wt/*R349P^ and wild-type tissues (Fig. 5A,C, Supplementary videos 1-3). While none of the wild-type tissues showed signs of rupture during or after tetanic stimulation, two of the four mutated tissues ruptured during tetanic stimulation. Furthermore, we observed rupture of individual fibers in the mutated tissues that survived tetanic stimulation (Supplementary videos 2-3).

**Figure 5.**
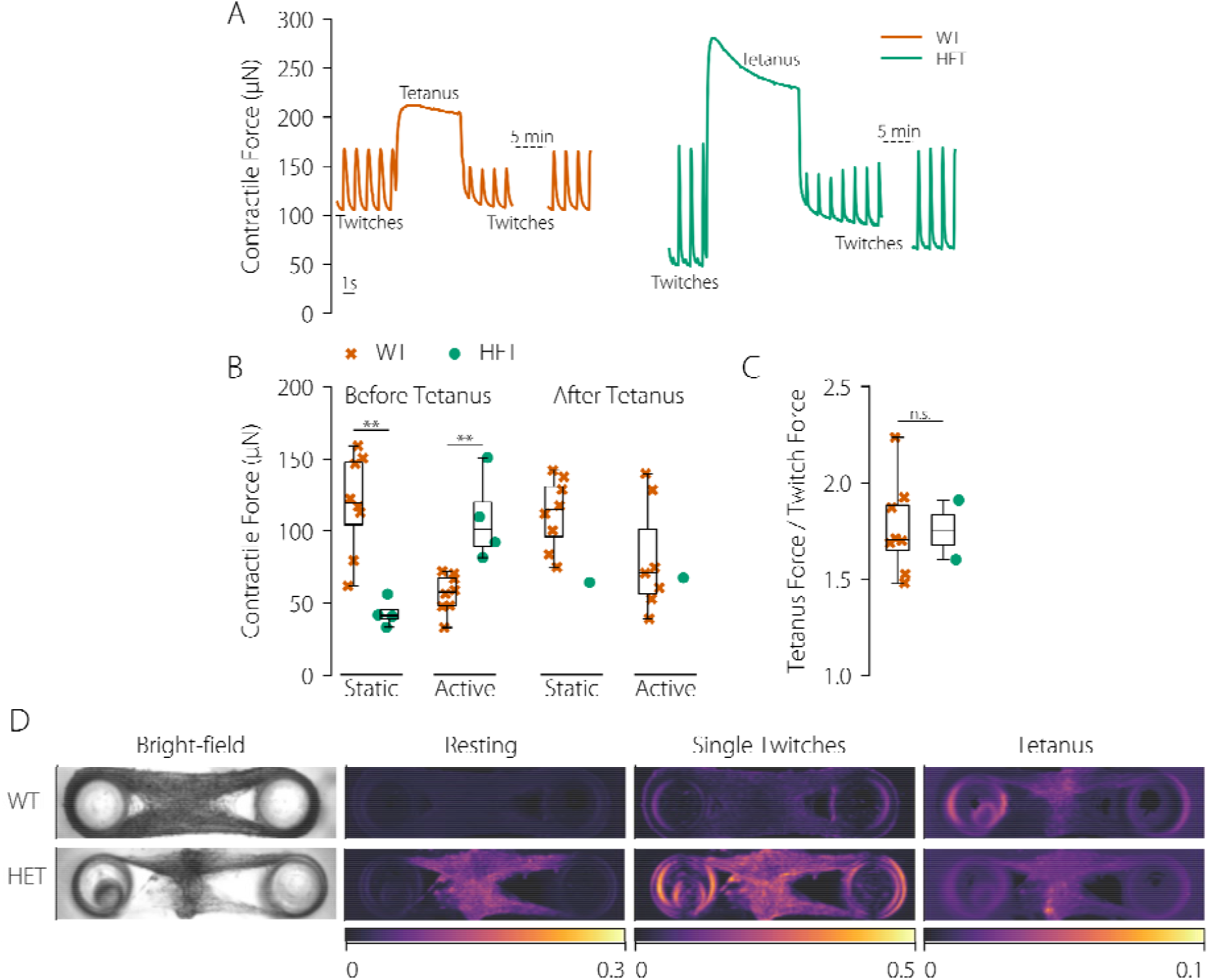
Comparison of wild-type and heterozygous desmin-mutated tissues. **A**. Representative force curve of wild-type tissue (orange) compared to Des^wt/R349P^ tissue (green) during single pulses and tetanic stimulation. **B**. Active and static contractile force (single pulse) of wild-type and heterozygous tissues before and 5 min after tetanic stimulation. Note that two of the four mutated tissues ruptured during tetanic stimulation. **C**. Ratio of maximum force increase during tetanic stimulation relative to active force during single pulse stimulation. **D**. Visualization of local contractile movements from the coefficient of variation in bright-field pixel intensities (left) during a resting period (5 s), a single twitch period (1 Hz frequency, 5 s), and during full tetanic activation (5 s) for a representative wild-type and Des^wt/R349P^ tissue. Scale bar 500 µm. *p ≤ 0.05, **p ≤ 0.01, ***p ≤ 0.001, n.s., not significant (p > 0.05).

Immediately following tetanic stimulation, single-twitch active forces were lower in both tissue types (Fig. 5A). Within 5 min of continuous single twitch stimulation, however, active forces of the wild-type tissues recovered, and in some tissues even increased beyond pre-tetanus levels (106 ± 79 µN after 5 min, compared to 57 ± 14 µN prior to tetanic stimulation). We could not test how active forces recovered in *Des*^*wt/*R349P^ tissues, as one of the two only tissues that survived after tetanic stimulation ruptured during the following single-twitch period. Static forces in wild-type tissues were not altered after tetanic stimulation (119 ± 35 µN before tetanic stimulation versus 112 ± 25 µN after 5 min; Fig. 5B). The single remaining *Des*^*wt/*R349P^ tissue, however, also showed no loss of static force. Together, these data suggest that the impaired integrity of *Des*^*wt/*R349P^ tissues and their increased tendency to rupture is caused by excessive mechanical tension induced by hypercontractility.

### Des^wt/R349P^ tissues show pronounced spontaneous, inhomogeneous contractions

In the absence of electrical stimulation, wild-type tissues were quiescent and did not display spontaneous contractions. In contrast, *Des*^*wt/*R349P^ tissues contracted spontaneously, even during the periods between two single pulses applied at 1 Hz (Supplementary video 2). These unsynchronized spontaneous contractions became particularly apparent in muscle fibers that were detached from the main tissues and accumulated in the central tissue regions. Because these muscle fibers do no longer experience any appreciable load, their spontaneous contractions result in large deformations. To visualize these local deformations, we calculated the coefficient of temporal variation (COV) of pixel intensities from bright-field image sequences recorded at 28 Hz during single twitch stimulation, tetanic stimulation, and during phases without electrical stimulation (“resting”) (Fig. 5D). A higher COV is represented in Fig. 5D with brighter colors and corresponds to larger contractile movements.

In the absence of electrical stimulation (resting phase), wild-type tissues displayed a coefficient of variation close to zero due to the lack of tissue movement, while for *Des*^*wt/*R349P^ tissues, the spontaneous contraction across the tissue caused irregular movements and thus, a high coefficient of variation. During single twitch stimulation (1Hz), wild-type tissues displayed a small coefficient of variation, indicating a homogenous contraction throughout the entire tissue, whereas *Des*^*wt/*R349P^ tissues showed a high coefficient of variation, demonstrating that the tissue strain was spatially highly variable, pointing to the presence of substantial shear forces between regions of different strain. During the plateau phase of tetanic stimulation, both wild-type and *Des*^*wt/*R349P^ tissues showed a low coefficient of variation and, hence, almost no movement as they remain in a maximally contracted state. These results indicate the presence of considerable spontaneous, unloaded contractile activity in *Des*^*wt/*R349P^ tissues.

## Discussion

In this study, we implemented an *in vitro* system for fabrication, long-term culture, and mechanical assessment of skeletal muscle micro-tissues. Micro-tissues were grown from primary satellite cells isolated from skeletal muscles of wild-type mice and mice carrying a heterozygous desmin R349P mutation.

Due to their regenerative stem-cell properties, satellite cells are ideal for skeletal tissue engineering (31). However, there are conflicting reports as to whether satellite cells can maintain their stem cell potency under culture conditions. Stem cell potency has been reported to decline within a few days after transferring isolated satellite cells into 2-D cell culture (32–34). By contrast, culturing satellite cells in 3-D micro-tissues resulted in the maintenance of a functional satellite cell subpopulation (as revealed by staining of Pax7) for at least 2-3 weeks, even though the cells had been expanded under 2-D culture conditions for up to five days after isolation (35).

In our study, we generated micro-tissues from cells that had been cultured and repeatedly passaged under 2-D conditions for considerably longer than five days before transferring them into a 3-D culture system. We found no decline in the contractile performance with higher passage numbers (tested in this study up to passage 14). Our observations are also in agreement with two recent studies that have reported the successful generation of micro-tissues from murine (36) and rat satellite cells (35). We have followed the protocol published in these studies for differentiating satellite cells into spontaneously contracting myotubes, which requires the addition of Matrigel to the collagen-I extracellular matrix. It is currently debated, which component or property of Matrigel is essential for this differentiation, but without Matrigel addition, micro-tissues were unable to generate active tension (unpublished).

When embedded in a dilute collagen-I/Matrigel biopolymer network, the satellite cells compacted the network within 24 hours of culture into micro-tissues that span between two elastic PDMS pillars. These pillars served both as mechanical anchor points and - through the measurement of pillar deflection - as force sensors. Geometrically, our setup is an upscaled version of a previously published assay for growing micro-tissues from murine skeletal muscle cells of the C2C12 cell line in a PDMS culture chamber fabricated using soft lithography (20). In contrast, our PDMS culture chambers are fabricated from a mold that has been machined with a conventional milling machine and drill press. Accordingly, our micro-tissues are longer by a factor of 4 (1.8 mm compared to 500 µm) and wider by a factor of 3-4 (300-400 µm compared to 100 µm) and are grown from 5,000 cells instead of 400 cells. There is no intrinsic advantage of this geometrical upscaling other than that it allows us to control the matrix density and number of seeded cells individually for each tissue.

Tissue formation was accompanied by the generation of static contractile forces that steadily increased over time and reached values between 50 and 100 µN within 2-4 days. After around four days in culture, the micro-tissues showed characteristics resembling native skeletal muscle, including myofibrillar striation and active contraction in response to electrical stimulation pulses when applied with a repetition rate of 1 Hz. When the repetition rate was increased to 100 Hz, micro-tissues developed a tetanic contraction that exceeded active single twitch forces by typically a factor of 1.7. Micro-tissues can be “skinned” (membrane-permeabilized) and respond to the increase in intracellular calcium with a force increase over a 10-fold range, with a half maximum effective Ca^2+^ concentration (EC_50_) of around 700 nM.

Taken together, our micro-tissues resemble the behavior of native muscle tissues, with several deviations. Most notably, the maximum tension of our micro-tissues is 2-3 orders of magnitude smaller than that of native murine muscle tissue. We estimated the maximum tension of wild-type micro-tissues as the sum of static and active forces normalized by the cross-sectional area, which we obtain from the width of the tissues in the mid-section and assuming a circular cross-section. Typical values of muscle tension are 675 ± 123 Pa (mean ± SD, n=8) during single twitch stimulation, and 871 ± 114 Pa during tetanic stimulation. Other studies reported slightly higher twitch tensions between 1-2 kPa for micro-tissues grown from C2C12 skeletal muscle cells (20, 21), and 1.7 kPa for micro-tissues grown from satellite cells (35). For comparison, intact *soleus* (SOL) and *extensor digitorum longus* (EDL) muscles of young/adult mice generate tetanic tensions of around 150-220 kPa (SOL) and 200-240 kPa (EDL), respectively (37–39).

Our micro-tissues contract against an auxotonic load, but the stiffness of the pillars is high in relation to the contractile force of the tissues so that the degree of shortening during a single twitch remains below 2%. In intact muscle, a shortening of 2% does not lead to an appreciable force reduction compared to isometric loading conditions (40). Contractile isometric twitch tension of single adult murine SOL and EDL fibers have been reported to be approximately 50 kPa and 100 kPa, respectively (41). By contrast, when primary isolated mouse *flexor digitorum brevis* (FDB) fibers are embedded in a soft 200 Pa (Young’s modulus) Matrigel, they generate low tension comparable to that of our micro-tissues (twitch tension 0.44 kPa at 1 Hz, tetanic tension 2.53 kPa at 100 Hz) (42). It is currently unknown whether this low tension in these primary FDB fibers is the result of excessive fiber shortening of more than 40% or due to other reasons such as cell adhesion to Matrigel.

Furthermore, micro-tissues show a reduced force-frequency relationship compared to primary skeletal muscle, for currently unknown reasons. At a pacing frequency of 100 Hz, the contractile force of the micro-tissues increased only by a factor of around 1.7, compared to a factor of 6.7 in adult SOL fibers, a factor of 3.8 in EDL fibers (37), and a factor of 5.7 in FDB fibers (42).

Micro-tissues generate static forces that are comparable in magnitude to the active force, whereas static forces (also called residual tension, or tonus) in isolated primary skeletal muscle fibers are largely absent. The origin of this high static force in micro-tissues is currently unclear. Static force develops in parallel to the collective compaction/remodeling of collagen and Matrigel fibers during the first 1-2 days after cell seeding and remains approximately constant thereafter; a behavior that has also been reported by others (19, 20). Static forces always develop before the micro-tissues gain the ability to actively contract in response to electrical excitation, and for that reason it has been suggested that the presence of static stress is a crucial factor for successful myotube differentiation and maturation (20).

In a sense, the term “static force” is a misnomer as it has also been actively generated by acto-myosin contractile tension. Because this tension is transmitted through extracellular matrix fibers, it is believed that matrix remodeling or matrix crosslinking can permanently “lock” the contractile deformation in place - akin to the one-way catch mechanism in a ratchet - and thus maintain tension also in the absence of actively generated forces. To which degree this really happens is debated, as both collagen and Matrigel display viscoelastic stress relaxation and creep (42–44) and thus cannot store elastic strain energy over prolonged time periods. Hence, the static tension within the ECM must at least occasionally be “renewed” by actively contracting cells. However, the term “static force” has become established in the literature, and we use it here to refer to the residual force measured in the absence of electrical stimulation, or to the minimum force between two twitches.

Despite their significantly lower contractile tension compared to primary skeletal muscle tissue, *in vitro* engineered micro-tissues offer several advantages. They provide quantitative readouts of muscle tissue mechanics and contractility over a period of several weeks under controlled conditions. In contrast, dissected muscles and single fibers can only be stored for a few days and, once mounted to force transducers, can only be used over a time period of several hours. Micro-tissue systems also offer higher measurement throughput, as many more samples can be obtained from a single animal, which reduces ethical concerns. The specific advantage of satellite cells as a cell source for micro-tissues is that they proliferate and differentiate into myocytes efficiently, and that they can be frozen and expanded in multiple passages (at least 14 passages in our hands) without loss of contractile performance. In addition, satellite cell-derived micro-tissues offer the possibility to study myocyte differentiation processes in a multicellular 3-D environment.

The main finding of this study is that micro-tissues derived from satellite cells with a heterozygous desmin R349P mutation, one of the most common mutations associated with desminopathies in humans, were prone to tissue-disintegration. This tissue disintegration recapitulates in part the progressive skeletal muscle degenerative processes in patients suffering from desminopathies (10). Notably, mutant micro-tissues did not exhibit any contractile impairment or weakness compared to control tissues; rather, they displayed hypercontractility that led to mechanical tissue disintegration. This finding raises the question whether skeletal myofiber hypercontractility is also involved in the progression of desmin R349P-related desminopathies in humans.

Desmin-mutated cells formed micro-tissues within 24 hours after cell seeding, similar to wild-type cells, but the shape of the desmin-mutated micro-tissues was considerably more variable in comparison. Subsequently, desmin-mutated cells started to delaminate from the peripheral matrix fibers and aggregated near the center of the micro-tissues. Around 72 hours after seeding, the matrix fibers near the pillars progressively thinned out, and micro-tissues started to rupture (completely detach from the supporting pillars). After one week in culture, all desmin-mutated micro-tissues had failed, whereas wild-type tissues were still intact, mirroring the loss of functional muscle over time in desminopathies.

Tetanic stimulation lasting for 5 s triggered the immediate rupture of more than half of the desmin-mutated micro-tissues. A video showing this process can be found in SI (video 3). The remaining desmin-mutated tissues showed at least some degree of structural damage after tetanic stimulation in the form of a partial rupture or detachment of cells and matrix fibers (SI video 2). As a consequence, the active force remained permanently reduced thereafter. No such rupture during tetanus occurred in wild-type tissues, and their active force fully recovered to pre-tetanus levels and beyond within 5 min.

The highly heterogeneous morphology of R349P desmin mutant tissues, the cell detachment and cell aggregation in the tissue center, and the progressive tissue rupture, can be potentially explained by three distinct mechanisms:

First, mutant cells may secrete more matrix-degrading proteases or secrete less matrix proteins so that the extracellular matrix in mutant tissues is mechanically more vulnerable and ruptures more easily when the myotubes begin to generate contractile tension. Such an altered secretory behavior of *Des*^*wt/*R349P^ cells, however, has previously not been reported, and this mechanism - although it cannot be excluded - remains speculative.

Second, desmin mutated cells may exhibit reduced adhesion strength to the matrix and, therefore, detach more readily when they contract. This mechanism, however, would leave intact matrix fibers behind, whereas we see that the matrix fibers progressively thin out as the cells recoil towards the tissue center. Moreover, magnetic tweezer measurements using fibronectin-coated beads on heterozygous murine *Des*^*wt/*R349P^ myoblasts and human *Des*^*wt/*R350P^ myoblasts revealed higher adhesion strength compared to their wild-type counterparts (45, 46). Therefore, we dismiss a reduced strength of individual ECM-adhesions *per se* as a potential mechanism.

However, reports in the literature suggest that the contractile forces in *Des*^*wt/*R349P^ myotubes couples to the ECM via dystrophin/dystroglycan complex- or integrin-based adhesions that are fewer and further apart (5, 47), leading to a reduced structural integrity of myofibrils and a non-uniform, uneven distribution of contractile stresses, increased local tractions, and shear deformations between parallel arranged but poorly connected myotubes. The notion that matrix fibers that are exposed to excessive tractions and shear forces locally rupture is also in agreement with the finding that matrix components and cells first start to disappear near the wedge-shaped bifurcation towards the pillars where the largest shear stresses can be expected. Consistent with a more uneven force distribution are furthermore reports that SOL and EDL fiber bundles from *Des*^*wt/*R349P^ mice disintegrate easily when strained (30, 45).

Third, desmin mutated myotubes may be able to generate higher contractile forces compared to wild-type myotubes, which would also lead to increased local tractions and matrix rupture. Support for this notion comes from our observation that the active forces of desmin mutated tissues are larger than those of wild-type tissues, even though the majority of the mutated cells are arranged in a disorganized clump near the tissue center where they cannot contribute to the measured pillar bending. Moreover, this larger force is generated by relatively thin bundles of muscle cells and matrix fibers and hence is transmitted across a much smaller cross section, leading to a much higher tensile stress. Desmin-mutated tissues show spontaneous twitching when they are not electrically paced, and they complete a single twitch in less than half the time that it takes wild-type tissue.

These signs of hypercontractility in desmin-mutated tissues are not seen, however, in whole isolated muscles from *Des*^*wt/*R349P^ mice (48). The authors of that study found no change in muscle mass-normalized twitch or tetanus force of EDL muscles compared to wild-type. Furthermore, the maximum force induced by calcium and caffeine stimulation in single SOL and EDL fibers and fiber bundles from *Des*^*wt/*R349P^ mutant mice showed no significant differences compared to wild-type (30, 45, 49). It is unclear why artificial micro-tissues grown from wild-type versus desmin-mutated satellite cells show these pronounced differences in their contractile behavior whereas *in-vivo*-differentiated muscle tissue and muscle fibers do not. It is conceivable that wild-type micro-tissues remain in an early state of differentiation, which would explain their low contractile tension, whereas desmin mutated tissues differentiate and mature more quickly. Another explanation could be that hypercontractile fibers do occur in desmin mutated tissue *in-vivo*, but they eventually rupture or detach from the matrix, are subsequently removed by tissue-resident macrophages, and hence escape detection.

Our observation that cell-generated contractile stresses during tetanic stimulation are sufficient to cause a mechanical disintegration of desmin-mutated micro-tissues is consistent with studies in desmin-mutated mice that link high-impact physical exercise to accelerated cardiac tissue damage (50, 51). Furthermore, the slow structural disintegration of desmin-mutated micro-tissues over time without noticeable loss in total force up until late disease stages mimics the clinical course of the disease in humans. Thus, we believe that bioartificial micro-tissues can contribute to a better understanding of the mechanisms that lead to desminopathies and possibly also other myopathies, and may in the future help to develop novel therapies.

## Author contributions

Conceptualization: RS, BF; methodology: MS, DK, CAD, WS, IT, BF; mouse model: CSC; satellite cell isolation: DH, SH; investigation: MS, DK, BR; software: DK, RCG, CAD; formal analysis: MS, DK, RCG, BF; writing original draft: MS, DK, IT, BF; supervision, admin, funding: OF, SH, RS, BF. All authors read, edited, and approved the final version of the manuscript text and figures.

## Data availability

The data of this study are available upon request from the corresponding author.

